# Zipper plot: visualizing transcriptional activity of genomic regions

**DOI:** 10.1101/077073

**Authors:** Francisco Avila Cobos, Jasper Anckaert, Pieter-Jan Volders, Dries Rombaut, Jo Vandesompele, Katleen De Preter, Pieter Mestdagh

**Affiliations:** Center for Medical Genetics, Ghent University, Ghent, Belgium.; Cancer Research Institute Ghent (CRIG), Ghent, Belgium.; Bioinformatics Institute Ghent from Nucleotides to Networks (BIG N2N), Ghent, Belgium.

## Abstract

**Summary:** Reconstructing transcript models from RNA-sequencing (RNA-seq) data and establishing these as independent transcriptional units can be a challenging task. The Zipper plot is an application that enables users to interrogate putative transcription start sites (TSSs) in relation to various features that are indicative for transcriptional activity. These features are obtained from publicly available datasets including CAGE-sequencing (CAGE-seq), ChIP-sequencing (ChIP-seq) for histone marks and DNase-sequencing (DNase-seq). The Zipper plot application requires three input fields (chromosome, genomic coordinate (hg19) of the TSS and strand) and generates a report that includes a detailed summary table, a Zipper plot and several statistics derived from this plot.

**Availability and Implementation:** The Zipper plot is implemented using the statistical programming language R and is freely available at http://zipperplot.cmgg.be

**Supplementary information:** Supplementary Methods available online.

## 1 Introduction

The introduction of RNA-sequencing (RNA-seq) has revolutionized the field of molecular biology, revealing that up to 75% of the human genome is actively transcribed (Djebali *et al.*, 2012). The majority of this transcriptome consists of so-called long non-coding RNAs (lncRNAs). Reconstructing accurate transcript models for these lncRNAs is a major challenge when processing RNA-seq data. In general, lncRNA transcripts are less abundant compared to protein coding genes (Derrien *et al.*, 2012), often resulting in a lack of junction reads from which transcript models are inferred. In addition, lncRNAs are frequently located in the vicinity of protein coding genes and could therefore represent unannotated extensions of untranslated regions (UTRs) rather than independent transcriptional units. Finally, transcript reconstruction from RNA-seq data often gives rise to large numbers of single-exon transcripts. Distinguishing single-exon fragments that represent independent transcriptional units from those that result from genomic DNA contamination or incomplete transcript assembly is not straightforward. These challenging putative lncRNA transcripts are frequently ignored or filtered out based on stringent conservation criteria (Chen *et al.*, 2016). Given the overall limited conservation of lncRNAs, such approach may result in a large number of false negatives. Alternatively, lncRNA transcript models can be refined and filtered by integrating complementary datasets on chromatin state (i.e. ChIP-seq for histone marks or DNase-seq) and transcript boundaries (i.e. CAGE-seq to mark the transcription start site (TSS) or 3P-seq to mark the 3’ end of polyadenylated transcripts) (Ulitsky *et al*., 2011). Transcripts for which the transcription start site coincides with a CAGE-peak and is in close proximity to a H3K4me3 or H3K27ac mark are more likely to be independent transcriptional units compared to transcripts that lack these features.

To study transcriptional independence we have created the Zipper plot, a visualization and analysis method that enables users to interrogate putative TSSs for association with CAGE-peaks, histone marks and DNaseseq peaks across a large collection of tissue and cell types. The method is available as a webtool and can help researchers to obtain more reliable lncRNA transcript models.

## 2 Implementation

ChIP-seq, DNase-seq and CAGE-seq data were obtained from various public repositories (Suppl. Figure S1A. See Online Supplemental Methods for detailed information) and processed to establish a peak-based database.

### 2.1 Database querying, Zipper plot construction and generation of html summary reports

The user can query the database by providing a list of genomic features (one per line), each consisting of three tab-separated fields: chromosome, genomic coordinate (hg19) of the TSS and strand (Fig. 1A). After selecting the data type of interest (i.e. CAGE, histone mark or DNase peaks), the user has the option to run the analysis in one sample type of interest or across all available sample types. In the first option, the user knows in advance in which tissue the set of genomic features are more likely to be expressed; with the second option, each individual genomic feature is analyzed across all samples and the sample in which the peak is most closely associated to the genomic feature will be retained for further analysis (Fig. 1B).

**Fig. 1.**
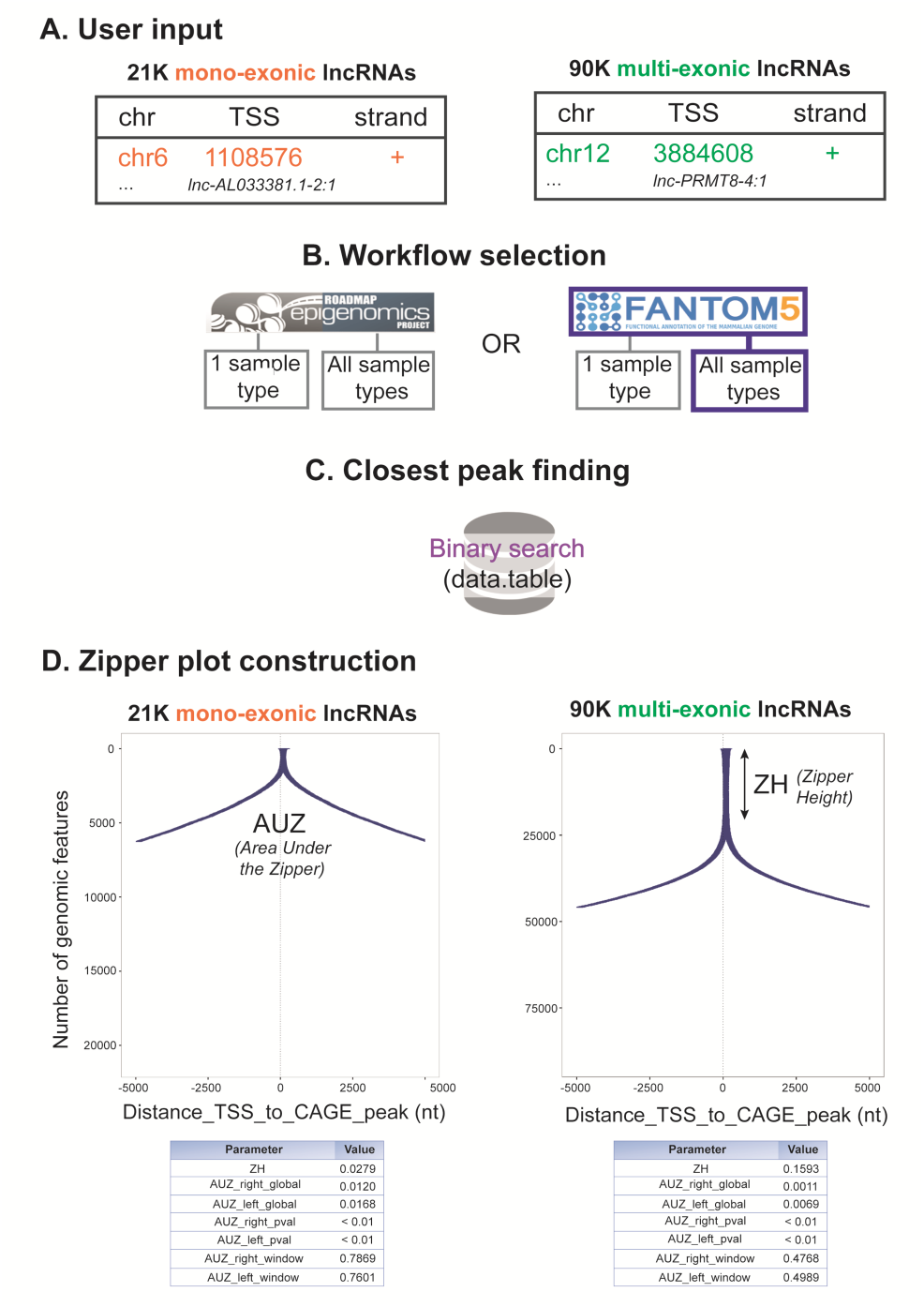
As an example application of the Zipper plot, we have studied all monoexonic versus all multi-exonic human lncRNA transcripts (21,102 and 90,508 respectively) from LNCipedia 3.1 (Volders *et al.*, 2015). We have selected the CAGE-seq data (FANTOM5) and “All sample types” workflow. As a result, 589 mono-exonic lncRNAs (2.8%) present a CAGE-peak overlapping with the TSS and 6,256 (29.7%) have a peak within a +/− 5kb window. On the other hand, 14.419 multi-exonic lncRNAs (15.9%) present a CAGE-peak overlapping with the TSS and 45.878 (50.7%) have a peak within a +/− 5kb window. These differences, also reflected in greater AUZ_global and AUZ_window values in the former case, might be reflecting the fact that numerous mono-exonic lncRNAs are not independent transcriptional units.

The data.table package (Dowle et al., 2015) is used to sort TSSs from the user’s input in a chromosome-wise manner and to perform a binary search (O(log n) time) in compiled C to retrieve the closest ChIP-seq/DNase-seq/CAGE-seq peak to each TSS (Fig. 1C).

Once this information is retrieved for all genomic features, the peaks are ordered based on the distance to the TSSs and a Zipper plot (for its similarity to an actual zipper) is generated with the aid of the ggplot2 package (Wickham, 2009) (Fig. 1D). Peaks overlapping with the TSS are placed at the top of the plot and the zipper starts to open as the peaks are located further away from the TSSs. By default the Zipper plot is visualized in a +/− 5 kilobase (kb) window around the TSS but the window size can be adjusted by the user. Suppl. Figure S1B shows in detail how the Zipper plot is built.

Two statistics named Zipper Height (ZH) and Area Under the Zipper (AUZ) are calculated for every Zipper plot. ZH corresponds to the quotient between the number of genomic features with a peak overlapping with the TSS and the total number of genomic features being studied (ZH ϵ [0,1]). The AUZ_global is computed as the sum of all the areas between the closest peak and the TSS of each genomic feature (Suppl. Figure S2. See Supplemental Methods). We have also incorporated the AUZ_window, which depends on the window size choice and is computed using only the retrieved peaks that lie within. This allows a quick visual comparison between two Zipper plots built using the same window size. Of note, ZH and AUZ are negatively correlated.

A one-sided p-value (AUZ_pval) is calculated by comparing the AUZ to those of random Zipper plots. The p-value represents the chance of finding a random Zipper plot with an AUZ_global smaller than or equal to the AUZ_global of the actual use case (see Online Supplemental Methods for more details).

Eventually, the knitr package (Xie et al., 2016) is used to generate an html report containing 1) the Zipper plot; 2) all the aforementioned parameters/statistics; 3) a summary table listing closest peaks, peak widths and overall peak enrichment information.

All the packages mentioned throughout this section are written in statistical programming language R.

## 3 Conclusion

The Zipper plot enables researchers to evaluate whether a set of putative lncRNAs have the characteristics of independent transcriptional units by integrating information on the 5’ boundary of the transcripts (i.e. CAGE-seq) and the chromatin state (i.e. ChIP-seq and DNase-seq) at the TSSs. More specifically, the presence of a CAGE-peak and activating histone mark at the TSS is indicative for independent transcription. Notably, the absence of such features does not imply the opposite. Low abundant transcripts may not show up in the CAGE-seq data because of too low sequencing depth or the expression of the lncRNA may be restricted to a tissue of cell type not included in the CAGE-seq, ChIP-seq of DNase-seq data.

Importantly, TSSs of RNA transcripts reconstructed from RNA-seq data might appear several nucleotides downstream of a CAGE-seq peak. Particularly for low abundant RNA transcripts, this inconsistency may be the result of an incomplete transcript assembly due to non-uniformity of read coverage towards 5’ ends and should be carefully examined.

To further expand the applicability of this tool, we plan to integrate publicly available data from methods that detect nascent RNAs (GRO-seq and PRO-seq), to extend the number of samples when new data becomes available and to allow users to work with their own data.

The Zipper plot is available at http://zipperplot.cmgg.be

## Acknowledgements

We thank Tom Sante for his technical advice and fruitful discussions throughout the development of this webtool.

